# The Geometry of Allostery: A Laplacian Minor Hierarchy for Many-Body Protein Communication

**DOI:** 10.64898/2026.06.10.731266

**Authors:** Fatma Senguler Ciftci, Burak Erman

## Abstract

Quantifying how cooperative, many-body relationships drive allostery in protein networks remains a major challenge. To address this, we develop the Laplacian minor hierarchy, a mathematical framework that characterizes the geometric invariants of a protein network. Lower-order minors yield standard metrics including the partition function and effective distances, whereas higher-order minors define novel topological measures: cooperation indices, each bounded between zero and one, that characterize pathway correlations at increasing levels of complexity, the third-order minor determines whether allosteric pathways are correlated or uncorrelated, and the fourth-order minor quantifies how distinct pathways communicate through intermediary residues. We apply this framework to analyze the evolutionary adaptation of the PSD95^pdz3^ domain from Class I to Class II ligand specificity via mutations G330T and H372A. The cooperation index demonstrates a distinct evolutionary hierarchy: the G330T mutation establishes distributed pathway couplings that the H372A mutation subsequently exploits, whereas H372A alone produces minimal global changes. Furthermore, the fourth-order analysis identifies His317 as a critical intermediary node bridging the class-switching (330–372) and class-bridging (330–400) allosteric pathways. These results demonstrate that allosteric dependencies emerge only when mutations accumulate in specific combinations, with a hierarchical organization of pathways structured around position 330 and intermediary nodes His317 and Phe400. Rather than predicting allosteric mechanisms, this framework provides a mechanistic explanation for why and how allostery emerges during protein evolution.

## 1. Introduction

Proteins execute their biological functions through the precise spatial arrangement of amino acid residues and the physical contacts established between residues that are distant along the primary sequence. Together, these contacts form a residue interaction network where amino acids act as interconnected nodes. The topology of this network dictates the protein’s mechanical stability, conformational flexibility, and intramolecular communication.

A local perturbation, such as a mutation, ligand binding event, or post-translational modification, propagates through this network to change the behavior of residues far from the point of origin. Understanding this phenomenon, known as allostery, requires quantitative metrics that reflect not just the simple geometric distance between two residues, but the full structural context of the pathways connecting them.

Standard structural analysis relies heavily on Cartesian or Euclidean distance. However, Euclidean distance is a function of only two points; it is entirely independent of the medium between them. While this abstraction is useful for rigid geometric measurements, it fails when applied to structured, cooperative biomolecules. In a protein, two residues separated by three bonds along a single solvent-exposed loop share a fundamentally different functional relationship than two residues separated by the same Euclidean distance but embedded within a highly packed hydrophobic core intersected by dozens of parallel interaction pathways. A metric that cannot distinguish between these two topological environments cannot fully capture allosteric communication.

The natural remedy is a distance metric that is not a property of two points in isolation but of their relationship within the full network, one that assigns shorter effective separation to residues connected by many parallel, high-weight pathways and longer effective separation to residues linked only by sparse or indirect routes [1–3]. This limitation is addressed by the concept of effective distance, which is derived from the generalized inverse of the graph Laplacian (a matrix representing network connectivity, detailed in the Theory section) [4–9]. Because the calculation of this metric fundamentally requires an algebraic inversion of the global connectivity matrix, the effective distance between any two points is intrinsically dependent on the entire network architecture. Every alternative pathway, parallel contact, and structural loop connecting all residues is simultaneously integrated into this single value. Because these effective distances satisfy the algebraic properties of inner products in a vector space, they endow the protein network with a genuine geometric structure in which every residue is represented as a displacement vector relative to a reference point, and distances between residues reflect their overall network connectivity rather than their spatial separation.

Network-based approaches built on related graph-theoretic foundations have made significant progress toward characterizing allosteric communication. By representing the protein as a graph, i.e., residues as nodes and interactions as weighted edges, spectral methods exploiting the eigenvalues of the graph Laplacian, contact network analyses [10, 11], and communication pathway algorithms have all demonstrated that network topology carries allosteric information invisible to Euclidean geometry [12, 13]. Yet these approaches share a common limitation: they are predominantly pairwise or global in nature, characterizing either the relationship between two specific residues or the overall spectral properties of the entire network. The many-body cooperative relationships among three, four, or more residues, the combinatorial fabric of allosteric sectors, remain largely inaccessible to these tools.

However, even effective distance itself shares this blind spot: it remains a pairwise metric. While the inversion of the Laplacian ensures that every interaction in the protein contributes to the final value, it still only calculates the communication strength between residue i and residue j. It cannot determine how a third residue k modulates or participates in that connection. In structural biology, allosteric mechanisms and cooperative transitions are rarely strictly pairwise; they frequently involve collective, multi-residue clusters on different paths but operating in unison, such as a catalytic triad or a coordinated hydrophobic pocket. To capture and quantify these higher-order collective interactions, a mathematical framework beyond pairwise metrics is required.

The mathematical structure needed to access this information is, however, already present in the Laplacian itself. It has long been established in graph theory that the minor hierarchy of the Laplacian matrix encodes the weighted spanning forest structure of a graph in a systematic and rigorous way, a result whose modern form is given by the all-minors theorem of Chaiken [14] and its extensions to weighted digraphs by Buslov [15]. In particular, for a selected residue set (S), the corresponding principal minor can be interpreted as a weighted sum over spanning forests whose components are rooted at the residues in (S). What has not previously been recognized is that this same hierarchy, when applied to the protein contact network, defines a natural sequence of residue-subset invariants that captures precisely the many-body cooperative relationships that pairwise and spectral methods cannot reach. The present paper develops this observation into a systematic framework, the Laplacian minor hierarchy, and demonstrates its application to allosteric communication in a model protein system, showing that successive minors provide geometric invariants of residue clusters that cleanly resolve the cooperative, competitive, and modular structure of the communication network.

Concretely, the successive minors correspond to familiar geometric and physical quantities:

a. The first minor counts the total number of weighted spanning trees of the contact network. By the Matrix-Tree theorem, this quantity is proportional to the partition function of the network, meaning that the full equilibrium statistical mechanics of the protein communication graph is encoded in this single scalar. Successive minors then build on this foundation as geometric invariants of residue subsets.
b. Second-order minors recover the standard pairwise effective distance (the length of a connectivity vector between two residues).
c. Third-order minors calculate the angle between two connectivity vectors relative to a shared reference residue, measuring the degree to which two distinct communication pathways overlap or diverge and defining a three-body interaction index.
d. Fourth-order and higher minors calculate multi-dimensional volume, quantifying the cooperative collective interactions within larger residue clusters that are mathematically invisible to purely pairwise measurements.

These geometric features, length, angle, and volume, are not arbitrary statistics imposed on the protein structure. Because all higher-order quantities derive directly from the pairwise effective distances, the hierarchy introduces no external parameters; each metric emerges as the natural mathematical consequence of expanding the graph Laplacian into its constituent minors, inheriting its physical meaning directly from the contact topology of the protein fold.

In this paper, we establish the mathematical foundations of this hierarchy and apply it to the well-characterized PDZ domain family. The first and second minors, corresponding to global network integrity and pairwise effective distance, respectively, have been developed in previous work [16, 17]. Here, we focus on the third-order minor and the three-body cooperation index it defines. We show that this index provides a direct topological readout of the allosteric state of the protein, tracking a known biological progression across ligand binding and successive mutations in a manner consistent with the functional epistasis characterized experimentally by Ranganathan and coworkers [18].

## 2. Methods

We represent the protein as a graph. Each residue is a node, and two residues are connected by an edge when they are close enough in three-dimensional space to interact. This is the same graph-based viewpoint used in Gaussian network models and related approaches to protein communication. Let *V* be the set of all residues, and let *N* be the number of residues. The graph is encoded by an *N* ⨯ *N* matrix called the Laplacian, denoted by *L*.

We construct the Laplacian matrix on the same basis as in the Gaussian Network Model (GNM) [10], employing a cutoff distance *r*_*c*_ such that only pairs of residues i and j with spatial separation *d*_*ij*_ ≤ *r*_*c*_ are considered as interacting, where *d*_*ij*_ denotes the Euclidean distance between the *C*_*α*_ atoms (or equivalent representative points) of residues *i* and *j*. The off-diagonal elements *L*_*ij*_ represent the interaction weights between residue pairs *i* and *j*, defined as:

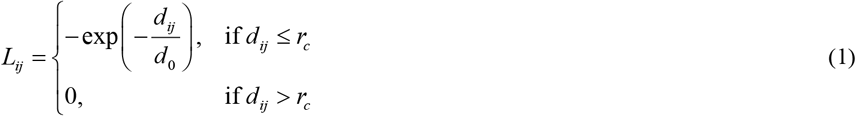

*d*_0_ is a characteristic length scale. This parameter *d*_0_ is interpreted as an effective temperature, since it can be written as *d*_0_ = *k*_*B*_*T* / *ε*, where *ε* is the constant energy per unit length (tension) of an effective linear potential *E*_*ij*_ = *εd*_*ij*_ governing the Boltzmann weighting. Earlier work [16, 17] showed that *d*_0_ = 1 is a reasonable value for the effective temperature, which is also used in the present work. The diagonal elements *L*_*ii*_ represent the total weighted connectivity of node *i*, ensuring the row-sum property of the Laplacian, 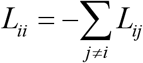.

By constructing the Laplacian matrix elements directly as exponentials of the scaled distances, the network’s topological invariants naturally map onto physical thermodynamic ensembles. Specifically, the first-order minor of *L* directly represents the partition function of the weighted spanning-tree ensemble, where the effective Boltzmann weight for an edge is exp (–*d*_*ij*_ / *d*_0_). This provides a rigorous statistical mechanical foundation for the subsequent metrics. [16, 17].

If *S* is a set of residues, *L* (*S*) means: delete from *L* all rows and columns whose labels are in *S*. For example, *L* (*i*) means delete row i and column i; *L* (*i, j*) means delete rows and columns i and j; *L* (*i, j,k*) means delete rows and columns i, j, and k. The symbol det means determinant. In this setting, determinants count network-wiring possibilities. Kirchhoff’s Matrix-Tree theorem says that det *L* (*i*) counts spanning trees, that is, the number of loop-free ways to connect the whole protein network. The all-minors Matrix-Tree theorem extends this idea to larger deleted sets *S* [14].

Throughout the steps below, i, j, k, l denote distinct residues.

Although *L* (*S*) is formed by deleting rows and columns mathematically, the biological meaning is not that the residues in *S* are physically removed. Rather, these residues are fixed as anchors. With one fixed residue, the determinant counts spanning trees of the network. With several fixed residues, it counts forests organized around those fixed residues. Thus, Laplacian minors describe how the protein network is organized around chosen reference residues.

### 2.1. Levels of the Minor Hierarchy

#### 2.1.1. Fixing one residue: total network integrity

Choose one residue i and form *L* (*i*). Define:

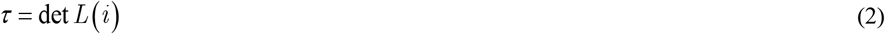

The number *τ* is the total number of spanning trees of the protein network, given by the determinant of any cofactor of the Laplacian matrix, as guaranteed by the matrix-tree theorem [16]. It does not depend on which residue i is deleted. This quantity has a direct connection to statistical mechanics: If the edge weights are interpreted as Boltzmann statistical weights, then *τ* is the partition function of the weighted spanning-tree ensemble. In that restricted ensemble, one may define an effective free energy *F*_*t ree*_ = –*k*_*B*_*Tlogτ*. This is not the full molecular Helmholtz free energy, but a graph-ensemble free energy associated with network connectivity.

In this sense *τ* encodes the total statistical weight of all configurations through which the network can transmit information, and its logarithm measures the thermodynamic cost of constraining the network to a single pathway. We use *τ* as the normalizing constant for all the quantities below. This normalization makes higher-order quantities easier to compare across proteins of different sizes, because they are measured relative to the total spanning-tree weight of their own network. In physical terms, normalizing by *τ* expresses each higher-order quantity as a ratio of statistical weights, analogous to computing a conditional probability within the ensemble of all network configurations. In other words, *τ* measures how many different ways the protein network can stay connected, weighted by the strength of each connection. A larger *τ* means more alternative routes are available, and equivalently, a lower free energy cost for information transmission through the network.

#### 2.1.2 Fixing two residues: pairwise effective distance

Now choose two residues i and j. Showing the second minor of the Laplacian by *L* (*i, j*) and using the matrix tree theorem [14]:

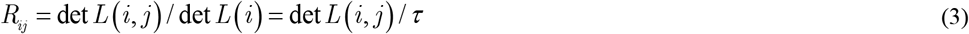

The number *R*_*ij*_ is the effective distance between residues i and j. It is the same mathematical object that graph theory calls effective resistance. Simple meaning: *R*_*ij*_ is not the physical distance in Angstroms. It is a network distance. If many paths connect i and j, then *R*_*ij*_ is small. If only a few paths connect them, then *R*_*ij*_ is large. Thus, *R*_*ij*_ is a generalized topological distance: smaller values mean that two residues are closer in the communication geometry of the protein network.

By Kirchhoff’s formula, the minor det *L* (*i, j*) and the normalization *τ* satisfy:

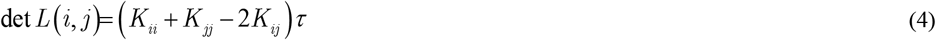

where *K* = *L*^+^ is the Moore-Penrose pseudoinverse of *L*. The effective distance can therefore be written equivalently as:

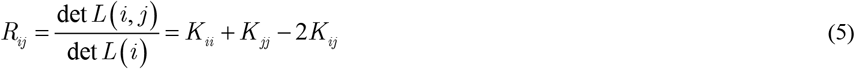

This expression is a sum of squared terms and is therefore never negative, equaling zero only when i=j. This mirrors the familiar Euclidean squared distance ‖*x* – *y* ‖^2^, with *L*^+^ playing the role of the metric. As shown in Appendix A, *K* defines a genuine inner product space in which every residue j is represented as a displacement vector *v*_*ij*_ from a fixed reference residue i, with squared length:

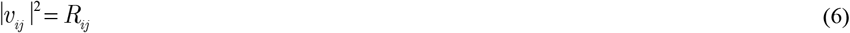

Residues that are tightly coupled through the contact network sit close to the reference in this space, while poorly connected residues sit far away. The choice of reference i sets the origin but does not affect the pairwise distances, since *R*_*ij*_ depends only on the network topology.

For two residues j and k, the inner product of their displacement vectors from reference i, denoted 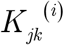, measures the degree to which the two communication directions from i toward j and from i toward k overlap in effective distance geometry:

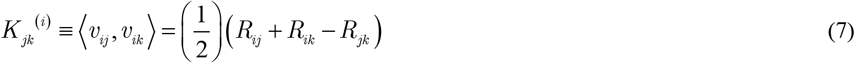

The derivation of Eq. 7 is given in Appendix B. The corresponding angle between the two communication directions is:

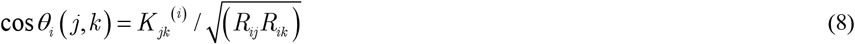

This is not a physical bond angle, it is a network angle. Small angles indicate that the two directions are strongly aligned in effective distance geometry, suggesting shared communication infrastructure. Angles close to 90° indicate nearly orthogonal directions, suggesting independent communication channels from the viewpoint of reference residue i.

A second important quantity is obtained by summing all effective distances from one residue i to the rest of the protein:

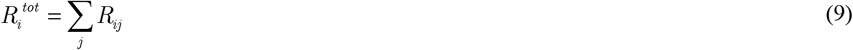

Here 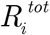 is the total effective distance of residue i. Small 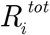 means residue i is globally central in the network. Large 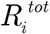 means residue i is more peripheral. We can also turn the row of distances starting from residue i into a probability distribution:

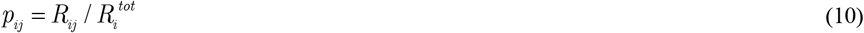

Thus, *p*_*ij*_ is the normalized share of the total effective-distance budget of residue i that is assigned to the pair i-j. In this sense, *p*_*ij*_ can be read as the probability weight of the i-j interaction in the row centered at residue i. Note: Since *R*_*ij*_ is a distance, large *p*_*ij*_ means that the pair contributes a large share of residue i’s total network distance.

The mathematical equivalence of the derivations in the Methods section to the Elastic Network Model (ENM) framework is detailed in Appendix D.

#### 2.1.3. Fixing three residues: triad coordination

Choose three residues *i, j*, and *k*. The pairwise effective distances *R*_*ij*_, *R*_*ik*_, and *R*_*jk*_ are defined by Eq 5, and the path-overlap 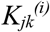 by Eq. 7. The triad effective distance is defined as:

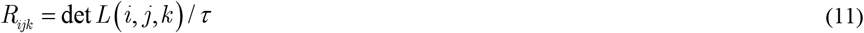

The Gram matrix of path overlaps relative to *i* is (See Appendix C for detailed explanation):

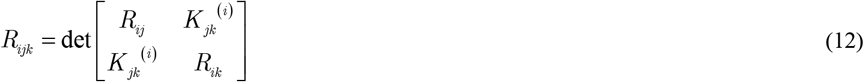

The number *R*_*ijk*_ is the effective distance of the three-residue set. It asks whether the three residues behave as separate pairwise relations or as a single coordinated unit. To compare the three-residue quantity with pairwise distances, use residue i as the reference point. Use Eq. 7 for the overlap between the two paths from i to j and i to k.

We define the cooperation index *χ*_*ijk*_, which quantifies the degree to which residue *i*’s pairwise couplings to *j* and *k* act as a coordinated unit rather than independently:

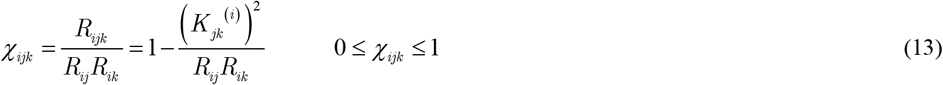

where the limits of *χ*_*ijk*_ are derived in Appendix C.

The Cooperation Index quantifies how a reference node i interfaces with the communication pathway between a pair of nodes j, k. This framework describes how the topological trajectories connecting i, j, and k intersect within the network graph. The two limits shown by Eq. 13 imply the following:

a. χ→1 (Low Pathway Overlap): The overlap term is small relative to the component distances, meaning the network trajectories from i to j and from i to k utilize largely separate structural components, i.e., distinct nodes and edges. The two communication channels are topologically disjoint and operate through independent subgraphs.
b. χ→0 (High Pathway Overlap): The overlap term is large relative to the component distances, indicating strong mechanical coupling. Perturbations at node i that propagate toward j and toward k are correlated; when one pathway responds to a perturbation, the other responds similarly, even if the two pathways involve different sets of residues.

#### 2.1.4. Fixing four residues: bond-bond interaction

A bond here means a pair of residues. Let one bond be (i, j) and another bond be (k, l). To study whether these two bonds are coupled, form *L* (*i, j,k,l*) and define:

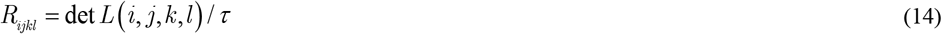

The number *R*_*ijkl*_ measures the joint effective distance of the two bonds. Using residue i as the reference, define for any two residues *a,b* ∈ {*j,k*,*l*} as placeholders:

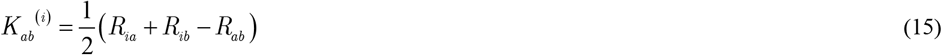

The effective distance yields a 3 x 3 determinant of path overlaps relative to the reference node:

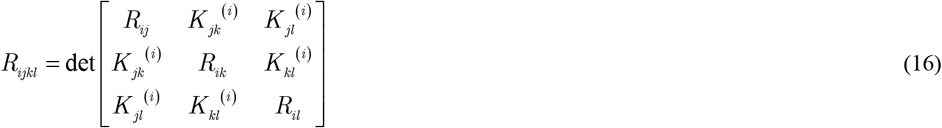

When the four-residue metric is lower than the baseline calculated from individual bonds, the two bonds exhibit cooperativity; conversely, a value higher than expected indicates competition. In this framework, a branch is defined as a bond extending from a designated residue. Consequently, by evaluating two specific branches, such as (*i, a*) and (*j, b*), the metric determines whether they interact cooperatively, competitively, or independently.

The fourth order cooperation index is similarly

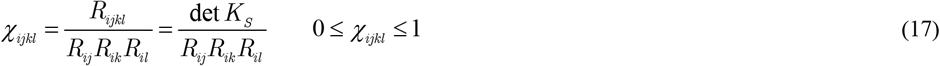

where the limits of *χ*_*ijk*_ are derived in Appendix C.

#### 2.1.5. Isolating any group of residues: the general rule

The first four steps are examples of one general formula. Let *S* = ?*s*_1_,*s*_2_,…,*s*_*r*_ ? be any group of r residues. Choose *s*_1_ as the reference residue. Define an (r−1) x (r−1) matrix *K*_*S*_ by:

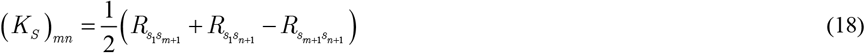

where m, n = 1, …, r−1. The effective distance of the whole group is:

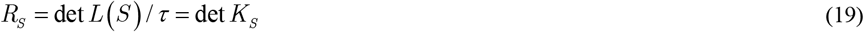

Simple meaning: Once all pairwise effective distances *R*_*ij*_ are known, all higher-order quantities can be computed from them. The pairwise distance table is like an alphabet. Triads, bond pairs, and domains are words written with that alphabet. Invariant viewpoint: Each minor gives an invariant of the constrained protein network. The first minor τ measures global network integrity through the total spanning-tree weight. Higher minors extend this idea to rooted forests and multi-residue constraints. The second minor gives the pairwise invariant *R*_*ij*_, describing dynamic separation and pairwise allosteric susceptibility. The third minor gives a triad invariant, measuring whether three residues behave independently or as a coordinated unit. The fourth minor gives a bond-bond, or branch-branch, invariant, measuring whether two local interactions cooperate or compete. Thus, the minor hierarchy assigns a characteristic network signature to each protein.

Details of the derivation of the general expression for the effective distance given by Eqs. 18 and 19 and its reduction to the second, third, and fourth order effective distance terms are outlined in Appendix C.

## 3. Comparison with experiments: Allostery of the PDZ domain

Ranganathan and colleagues [18] mapped the evolutionary transition of a PDZ domain from Class I to Class II ligand specificity using PSD95^pdz3^. The wild-type PDZ binds Class I ligands (e.g., CRIPT) but does not bind Class II ligands (e.g., T-2F) due to steric clash between the bulky phenylalanine at the −2 position and H372. H372A removes this histidine, creating space for the phenylalanine; the mutant gains partial T-2F binding but loses Class I affinity. G330T, an allosteric mutation on the β2–β3 loop, stabilizes an alternative conformation; alone, it enables high-affinity T-2F binding while preserving Class I recognition, a dual-specificity intermediate. Combining G330T with H372A yields complete Class II specialization: the double mutant binds T-2F with high affinity and no longer binds Class I. Thus, G330T is conditionally neutral; it enables T-2F recognition without compromising Class I binding upon a change in conditions.

### 3.1 Mapping cooperative path dependence onto the PDZ allosteric landscape

We computed the cooperation index *χ*_*ijk*_ and its mutation-induced changes Δ*χ*_*ijk*_ = *χ*_*ijk*_ (*Mutated*) – *χ*_*ijk*_ (*WT*) across the four PSD95^pdz3^ variants (WT, G330T, H372A, and G330T+H372A) in the T-2F ligand-bound state using crystal structures from the Protein Data Bank (PDB IDs: 5HED, 5HF1, 5HFC, 5HFF, respectively).

Figure 1 displays two heatmap panels with residue indices on both axes, showing only residues with |Δ*χ*| > 0.1 (greater than 10 percentage point increase in cooperativity) are shown. We computed the three-body cooperation index for all possible residue triplets; while this calculation is computationally efficient (< 1 minute on a desktop computer), visualizing results across three free variables is impractical. Since G330T is the primary allosteric mutation in this system, we fixed i=330 as the reference node, and systematically computed Δ*χ*_330, *j*,*k*_ to identify pathways containing residues j and k emanating from this central node. Only the*C*_*α*_ atoms were considered, and the cutoff distance between alpha carbon pairs in forming the Laplacian was set to 7.8Å for all structures. All calculations were performed according to the procedure given in [17].

**Figure 1.**
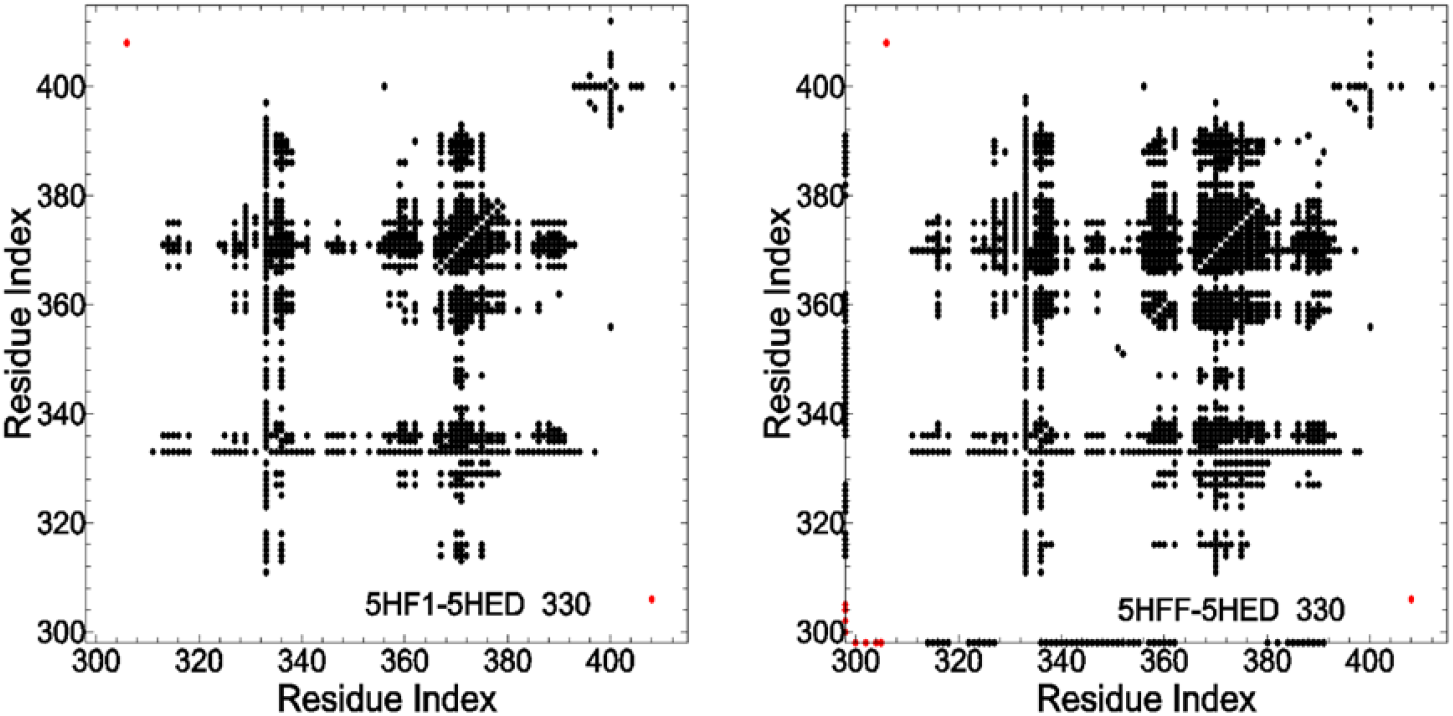
Global pattern of pairwise correlations in conformational changes by computing Δ *χ*_330, *j*,*k*_ values for i=330, between each variant and wild-type. Only residues with Δ*χ* > 0.1 are shown: Left panel (G330T vs WT, T-2F ligand) shows the Δ*χ*_330, *j*,*k*_ values between G330T and wild-type. Most values are negative, indicating coordinated conformational changes throughout the protein structure in response to the G330T mutation. Right panel (G330T,H372A vs WT, T-2F ligand) shows the Δ*χ*_330, *j*,*k*_ values for the double mutant. The right panel displays substantially more points than the left panel, indicating that the additional H372A mutation introduces significantly more correlations into the network of conformational pathways. This expansion of coupled residue pairs demonstrates that pathway interdependence increases dramatically when both mutations are present.

The left panel compares G330T (5HF1) versus WT (5HED), both with T-2F bound. The right panel compares the double mutant G330T+H372A (5HFF) versus WT, both with T-2F bound. Nearly all points in both panels show negative Δ*χ*_330, *j*,*k*_ values, indicating that both mutations enhance conformational coupling between residue pairs relative to WT. The right panel contains more points than the left, demonstrating that adding the H372A mutation to the G330T background introduces additional correlations into the network paths. This agrees with Ranganathan’s finding that the double mutant achieves a distinct, specialized Class II binding state, requiring a reorganized cooperative network.

The proliferation of correlation points in the double mutant compared to the single G330T mutant shows that H372A does not act in isolation; rather, it becomes embedded within a landscape of pre-existing conformational couplings established by G330T, creating an integrated allosteric network.

We next examined how the cooperation index varies across the protein sequence by computing Δ*χ*_330,372,*k*_ in the T-2F ligand-bound state. This triple specifically measures how the conformational relationship between the class-bridging site (330) and the class-switching site (372) is modulated by each residue k throughout the protein. Figure 2 plots Δ*χ*_330,372,*k*_ versus residue index (residues k=298–415) for three mutations, all in the T-2F-bound state. The green curve (H372A alone) shows Δ*χ*_330,372,*k*_ values near zero (order of 0.01), indicating that this active-site mutation introduces almost no change in cooperativity across the network. The black curve (G330T alone) shows significant negative Δ*χ*values, indicating that this distal allosteric mutation broadly enhances cooperativity between positions 330, 372, and residues throughout the protein. The red curve (G330T+H372A) shows Δ*χ*values that are more negative than the black curve, but not dramatically so. These findings agree with Ranganathan’s experiments: G330T is the key mutation that remodels the cooperative architecture to enable Class II recognition, while H372A alone does little to alter global cooperativity. The double mutant further reinforces this remodeling, consistent with its complete specialization to Class II binding.

**Figure 2.**
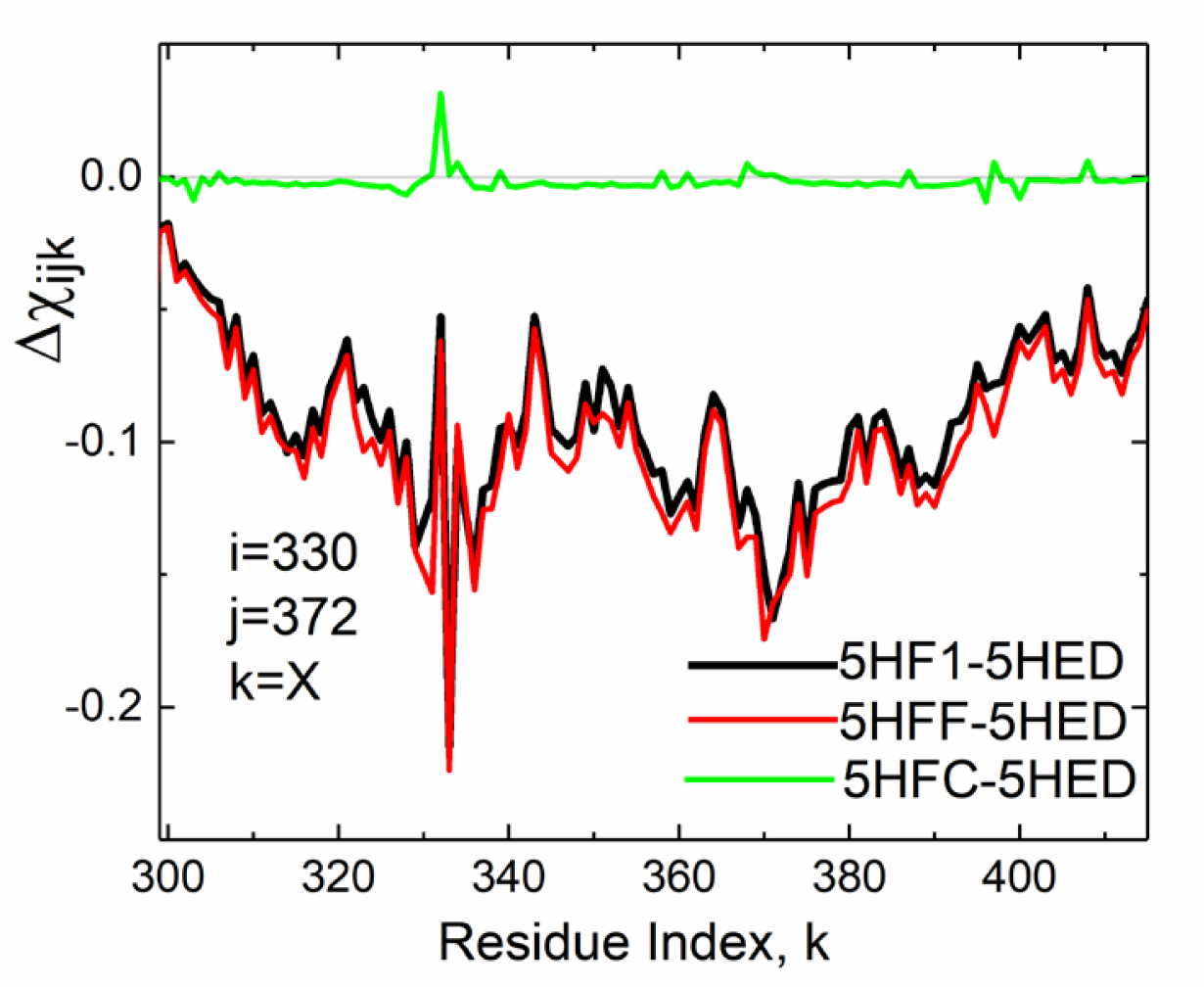
How the cooperation index varies across the protein sequence. The black curve (G330T mutation) shows significant negative Δ*χ*_330,372,*k*_ values across multiple positions, indicating that G330T creates measurable conformational coupling between positions 330, 372, and distant residue positions throughout the protein. The pattern reflects allosteric perturbations propagating from the β2– β3 loop through the protein structure. The green curve (H372A mutation alone) shows Δ*χ*_330,372,*k*_ values near zero (order of 0.01), indicating that H372A produces almost no detectable change in the conformational relationships between 330, 372, and other residues. This demonstrates that the H372A class-switching mutation, while effective at altering ligand specificity through direct active site perturbation, does not create the distributed pathway couplings characteristic of allosteric mechanisms. The red curve (G330T, H372A double mutant) shows negative Δ*χ*_330,372,*k*_ values more strongly negative than the black curve across most positions. The enhanced cooperation in the double mutant indicates that H372A amplifies and extends the allosteric network established by G330T. The additional mutation does not operate in isolation but becomes coupled to the pre-existing conformational framework, resulting in a more extensively interconnected network of cooperative residue interactions.

Figure 3 plots the fourth-order effective distances R(330, 372, 400, x) versus residue index (residues k=298–415) for three mutations, all in the T-2F-bound state. The fourth-order minor *R*_*ijkl*_ measures the joint effective distance of four residues, capturing whether two distinct pathways interact cooperatively (small R indicating coupling) or competitively/independently (R values closer to one indicating decoupling).

**Figure 3.**
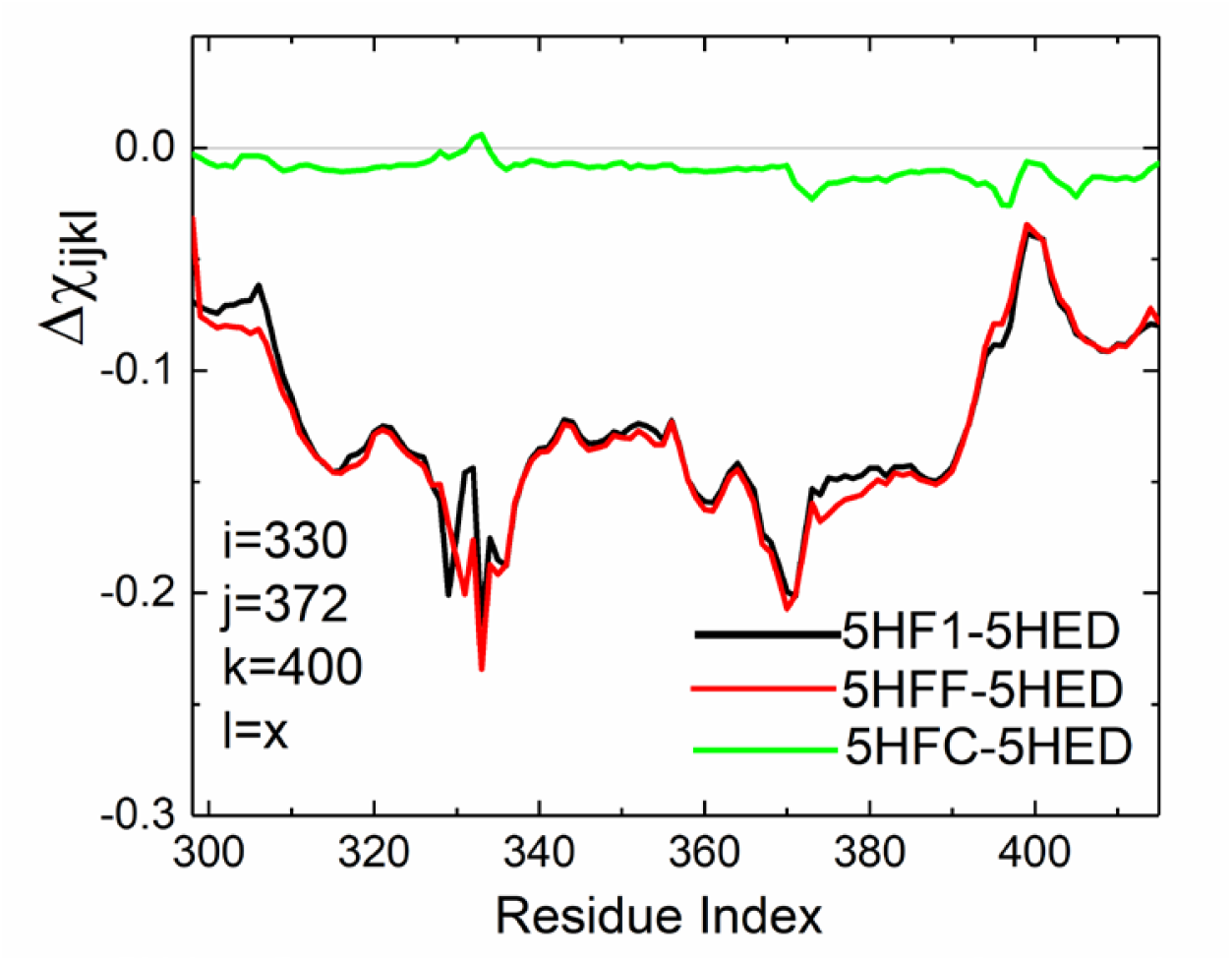
Fourth-Order Network Analysis Identifies Critical Intermediary Nodes in Allosteric Path Coupling. Fourth-order effective distance R(330, 372, 400, x) plotted against residue index x (residues 298–415) for three variants, each compared to wild type (PDB 5HED): G330T (black, PDB 5HF1), H372A (green, PDB 5HFC), and G330T+H372A (red, PDB 5HFF). Minima identify residues x that couple the 330–372 and 330–400 branches into a compact four-body unit. See legend for Figure 2.

The three curves represent: Black line, G330T (PDB 5HF1); Red line, G330T+H372A (PDB 5HFF); Green line, H372A (PDB 5HFC), all compared to WT (PDB 5HED). The analysis focuses on the quartet (330, 372, 400, x), where 330 is the allosteric hub, 372 is the class-switching active site, 400 is a known allosteric position showing reduced cooperativity in the heatmaps of Figure 1, and x varies across the entire protein.

WT, G330T and G330T+H372A show minima at positions 317 (His317), 330, 372 and 400–410. Residues 330, 372 and 400 are spatially proximate and define the core allosteric triangle; His317, by contrast, is spatially distant from this group yet consistently appears as a minimum of the fourth-order cooperation index. This is precisely what the framework identifies as a relay residue: His317 is not part of the local structural rearrangement centered on 330, but its fluctuations are simultaneously correlated with both functional branches 330–372 and 330–400, making the four-body interaction more compact than the two branches would predict independently. In other words, His317 couples the two branches into a single coordinated unit without being physically adjacent to either endpoint. The minimum at His317 is present in WT, G330T and G330T+H372A, and a less pronounced feature at the same position is also discernible in H372A. That this position is consistently identified across all variants, including the one with the weakest allosteric reorganization, suggests that His317’s relay function reflects a conserved topological feature of the PDZ fold rather than a mutation-induced effect. His317 therefore represents a fixed architectural element of the allosteric network, a residue whose intermediary role is encoded in the fold itself rather than induced by either mutation. Equally notable is that H372A alone leaves the fourth-order cooperation index unchanged, despite its decisive functional effect on ligand specificity. This indicates that H372A acts as a local gating residue, it removes the steric barrier to Class II binding without reorganizing the cooperative path structure of the network. That H372A produces only a weak cooperative signal overall, without shifting or removing the minimum at H317, is consistent with its role as a local gating residue whose effect is confined to the active site. The allosteric redistribution of mechanical coupling is carried entirely by G330T; H372A is a local enabler of the conformational change that G330T prepares. The fourth-order framework thus draws a mechanistic distinction between the two mutations that binding affinity data alone cannot provide.

### 3.2 Interpretation: Emergent Pathway Interdependence and Hierarchical Network Organization

The three curves in Figure 2 show the mechanistic basis for why G330T, H372A succeeds evolutionarily, where H372A alone does not. H372A (green curve) acts through direct, localized perturbation of the active site, with minimal effect on global network cooperativity. It achieves partial class switching through a steric mechanism but leaves the binding pocket structurally rigid and unable to accommodate both ligand classes simultaneously. G330T (black curve) establishes an allosteric backbone, a distributed network of pathway couplings throughout the protein structure, even though G330T itself makes no direct ligand contact. This allosteric effect is the signature of a network-level reorganization that increases conformational plasticity at the binding pocket. G330T, H372A (red curve) does not simply superimpose independent effects; rather, H372A operates within the conformational landscape prepared by G330T, resulting in enhanced and extended pathway interdependence across the entire protein.

This pattern demonstrates that the cooperation index Δχ directly shows emergent pathway dependencies that arise only when mutations accumulate in specific combinations. The expansion of negative Δχ values in the double mutant compared to either single mutant shows that allosteric coupling is not an inherent property of individual mutations, but rather an emergent property of the evolved network, a signature of functional integration that arises when the system reorganizes to accommodate both mutations simultaneously. In evolutionary terms, G330T creates the allosteric capacity; H372A leverages it.

The fourth-order analysis in Figure 3 shows the hierarchical organization underlying this emergent cooperativity. The identification of His317 as a critical intermediary node shows that the allosteric network is not a random distribution of couplings but a structured hierarchy centered on position 330. His317 acts as a bottleneck, a control point through which the class-switching (330–372) and class-bridging (330–400) pathways are integrated. The invariance of this minimum across all variants demonstrates that His317’s role as a topological relay is a conserved property of the PDZ fold, not an emergent consequence of either mutation. The network’s hierarchical organization through His317 is therefore a pre-existing architectural feature that both G330T and H372A use, rather than one they create. Mechanistically, this suggests that allosteric communication does not flow through all possible routes with equal probability. Instead, the network exhibits preferred pathways that are fixed in the fold: perturbations originating at position 330 reach and coordinate the 372 and 400 branches most efficiently by routing through His317, regardless of the mutational background.

The fourth-order insight refines our understanding of the three-body cooperation index. While Δ*χ*_330,372,400_ tells us whether the two branches are coupled, R(330, 372, 400, x) tells us how that coupling is implemented, through which intermediary residues the network traffic flows. This nested hierarchy, from three-body cooperation indices to four-body effective distances, demonstrates that the Laplacian minor framework moves beyond a binary yes/no question about cooperativity to a precise identification of the residues through which cooperativity is structurally organized. That this intermediary, His317, is fixed across all mutational backgrounds while the coupling strengths at 372 and 400 respond to mutation suggests a clean division of labor: the fold provides the routing architecture, and the mutations modulate the signal carried through it.

## 4. Discussion

This study demonstrates that the Laplacian minor hierarchy provides a powerful framework for understanding allosteric mechanisms at the level of many-body cooperativity. By computing successive minors of the protein contact network, we can quantify not only pairwise effective distances but also three-body cooperation indices and four-body branch-coupling metrics that show the hierarchical organization of allosteric pathways.

Our analysis of the PSD95^pdz3^ evolutionary adaptation provides clear evidence that allostery is not a property of isolated mutations but emerges from network reorganization. This is a manifestation of epistasis at the network level: G330T, located distant from the ligand-binding site, establishes distributed pathway couplings that H372A subsequently exploits. H372A alone cannot achieve complete class switching because it operates within a rigid network; combined with G330T, it gains the conformational freedom necessary for Class II specialization.

This framework provides the structural and mechanical basis for these evolutionary paths, directly complementing sequence-level observations. While Ranganathan and colleagues established the presence of an allosteric sector in PSD95^pdz3^ through sequence-based Statistical Coupling Analysis (SCA) [18], SCA inherently identifies co-evolving residues without mapping the physical, multi-body pathways through which they communicate structurally. Our Laplacian minor hierarchy bridges this mechanistic gap. By evaluating the network topology via higher-order invariants, the cooperation index gives structural form to functional epistasis: it shows how the active-site H372A mutation physically utilizes a pre-reorganized topological landscape established by G330T, routed through specific structural intermediaries like His317. Consequently, this topological mapping clarifies the nature of conditional neutrality at the level of network topology:

G330T is neutral in the Class I binding environment but becomes essential for H372A’s function in the Class II environment. The fourth-order analysis showing His317 as a critical intermediary node provides a concrete mechanistic explanation for this functional dependency, showing how allosteric epistasis arises from hierarchical network organization rather than simple pairwise interactions.

His317 is not a residue involved in direct ligand binding or major conformational change; rather, it serves as a network hub through which allosteric information flows. This finding suggests that allosteric mechanisms can be understood not merely as long-range conformational propagation but as a hierarchical routing of information through structured intermediary nodes.

While this study focuses on the PDZ domain, the framework should generalize to other allosteric systems. The Laplacian minor hierarchy makes no assumptions about the specific protein; it depends only on network topology, which can be computed from any crystal structure. Future studies could test whether:

1. Other allosteric proteins show similarly structured hierarchies of intermediary nodes;
2. Evolution frequently optimizes allosteric mechanisms by concentrating control through a limited set of strategic intermediaries;
3. The fourth-order and higher minors show vulnerabilities or robustness in allosteric networks, with implications for disease-causing mutations.

The framework could also be extended to incorporate dynamic information from molecular dynamics simulations. The current analysis uses equilibrium structures; weighted versions of the Laplacian incorporating residue contact frequencies or correlation matrices from MD trajectories could show how allosteric hierarchies shift between conformational substates.

From a theoretical perspective, the Laplacian minor hierarchy connects network topology to functional phenotype in a way that previous approaches could not. Pairwise effective distances show which residues are globally central and which are peripheral; the cooperation index shows which pairs of pathways are integrated or independent; the fourth-order minor shows the critical nodes structuring that integration. Each level of the hierarchy adds mechanistic resolution without requiring external parameters, every quantity derives from the contact network alone.

This hierarchical approach also offers a fresh perspective on why allostery might have evolved. If allosteric regulation is expensive (requiring the protein to maintain distributed pathway networks), why do proteins possess it? Our results suggest that allostery emerges as a byproduct of evolutionary adaptation: proteins that can readily access new functions through mutations that create distributed pathway couplings, i.e., the conditional neutral mutations emphasized by Ranganathan, will be selected for adaptability even if the allosteric machinery is not initially favored for regulatory function. Over time, once these networks exist, they become coopted for functional regulation. This suggests that the origins of allostery lie in evolvability, not in function. This hypothesis is consistent with the observation that allosteric mechanisms appear universal in proteins that must adapt to changing environmental conditions.

Several limitations should be noted. First, the analysis assumes that the protein contact network is the primary pathway for allosteric communication. While this has strong empirical support from Gaussian network models and other approaches, alternative pathways (electrostatic fields, hydration dynamics) may also contribute. Second, we use static structures rather than dynamics; the hierarchies identified here are averaged over equilibrium fluctuations but do not capture transient interactions. Third, the fourth-order analysis depends on identifying and computing *R*_*ijkl*_ for relevant quartets; this requires prior knowledge of which quartet to examine. Future work should develop systematic approaches to identify critical quartets without prior bias.

## 5. Conclusion

We have established the Laplacian minor hierarchy as a rigorous, parameter-free framework for quantifying many-body cooperativity in proteins. Applied to the PDZ domain’s evolutionary adaptation to new ligand specificity, this framework shows that allostery emerges as a hierarchical network phenomenon where distributed pathway couplings (captured by the three-body cooperation index) are organized around critical intermediary nodes (captured by the fourth-order effective distance). These results suggest a mechanistic answer to a fundamental question: not “does allostery exist?” but “how is allosteric organization structured within the protein contact network, and what evolutionary pressures drive the emergence of these structures?” The answer appears to be that evolution selects for proteins capable of rapid adaptation through mutations that create pathway interdependence, and these adaptive mechanisms subsequently become repurposed for allosteric regulation of function. This perspective unifies evolution and mechanism, offering a framework for understanding how proteins satisfy the competing demands of structural stability and functional adaptability.

## Conflict of interest

The authors declare no conflict of interest.

### Data availability statement

All source code and data preprocessing scripts are publicly available at https://github.com/fatmasenguler/KRAS_spanning_tree

### Funding

This research received no external funding.

## Appendix A: The Inner Product Space of Effective Distances

We show that the pseudoinverse *K* = *L*^+?^defines a genuine inner product space in which the effective distances *R*_*ij*_ are squared distances between residues.

The graph Laplacian *L* is symmetric and positive semidefinite with a one-dimensional null space spanned by the constant vector 1. Its Moore-Penrose pseudoinverse *K* = *L*^+?^exists uniquely on the subspace orthogonal to 1 and inherits the symmetry and positive semidefiniteness of *L*. The spectral expansion of *K* is:

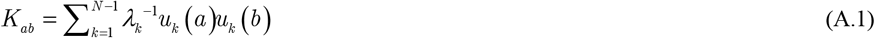

where the sum runs over the *N* –1 non-zero eigenvalues *λ*_*k*_ of *L* and *u*_*k*_ are the corresponding orthonormal eigenvectors.

Fix a reference residue i and define the displacement vector of residue j relative to i as:

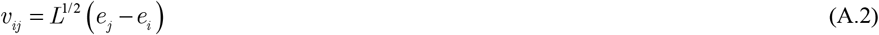

where *L*^1/2^ is the positive semidefinite square root of *K*, defined by *L*^1/2^*L*^1/2^ = *K*, and *e*_*j*_ is the standard basis vector with a one in position j and zeros elsewhere. The inner product between any two displacement vectors is then:

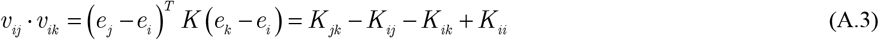

Setting k=j gives the squared length:

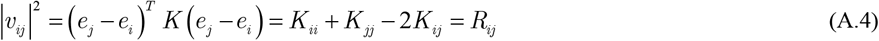

which recovers the effective distance exactly. Since *K* is positive semidefinite, |*v*_*ij*_|^2^≥0 with equality only when i=j, confirming that *R*_*ij*_ is a valid squared distance. The effective distance space is therefore a well-defined inner product space in which each residue is a point, the squared distance between any two points is *R*_*ij*_, and the inner product between displacement vectors is given by Eq. A.3. The choice of reference residue i sets the origin of the space but does not affect any pairwise distance, since *R*_*ij*_ depends only on the network topology through *K*.

## Appendix B: The Inner Product 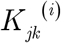 in Terms of Effective Distance

We derive the expression for the inner product 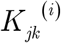 entirely in terms of pairwise effective distances, showing that no quantities beyond the second-order minor are needed to compute the three-residue geometry.

The effective distance between any pair of residues is defined from the pseudoinverse *K* = *L*^+^

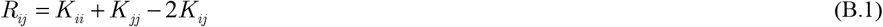

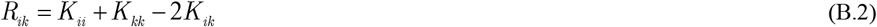

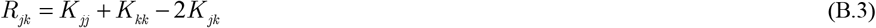

We solve each equation for the off-diagonal element of *K* :

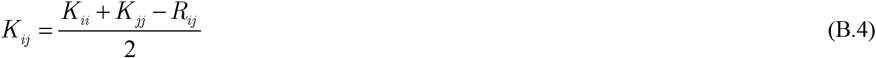

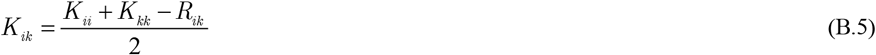

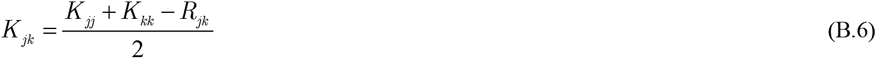

From the definition of the displacement vector *v*_*ij*_ = *δ R*_*j*_ – *δ R*_*i*_ and the inner product structure of *K*, the overlap between the two paths rooted at i is:

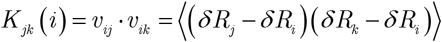

Expanding:

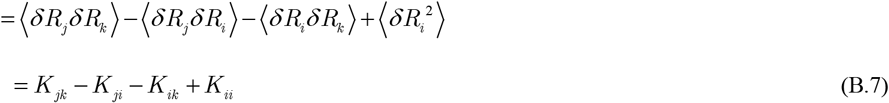

Substituting equations (B.4), (B.5), and (B.6) into (B.7):

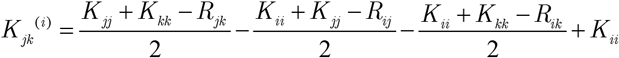

Expanding all terms:

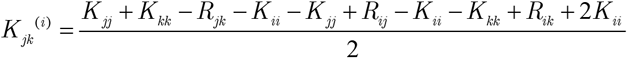

What remains is:

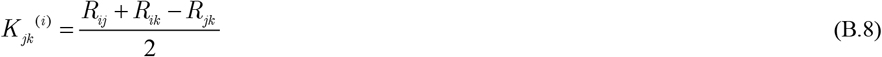

This is the distance analog of the law of cosines. The diagonal elements of *K* which depend on the arbitrary choice of gauge in the pseudoinverse cancel completely, leaving a result that depends only on the three pairwise effective distances. This cancellation is not accidental: it reflects the fact that effective distances, and therefore all higher-order quantities in the minor hierarchy, are gauge-invariant quantities independent of the arbitrary constant that can be added to the diagonal of *K*.

As an immediate consequence, the squared normalized overlap between the two paths rooted at i is

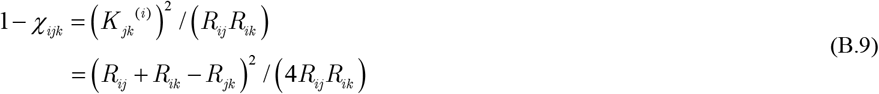

Eq. B.9 reduces to the cooperation index expression used in the main text.

Thus, the three-body cooperation index is expressed entirely in terms of the three pairwise effective distances of the triad. Once the pairwise effective distances R_ij_, R_ik_, and R_jk_ are known, the third-order normalized minor χ_ijk_ can be computed algebraically.

## Appendix C: The General *m*-th Order Effective Distance and Cooperation Index

In the Methods section, the effective distance of an arbitrary residue set was defined by a Laplacian minor. Here we prove that this definition is equivalently the determinant of an inner-product matrix built only from pairwise effective distances. We then define the normalized all-order cooperation index and prove that it lies in the interval [0,1]. Throughout, the graph is connected, so the Laplacian *L* is symmetric positive semidefinite and has a one-dimensional null space spanned by the constant vector **1**. The pairwise distances *R*_*ab*_ are effective distances [19]. The determinant identity below uses Jacobi’s complementary-minor identity [20], the combinatorial interpretation uses the all-minors matrix-tree theorem [14], and the range proof for the normalized cooperation index uses Hadamard’s determinant inequality for positive semidefinite Gram matrices [20].

### C.1 Statement

Let

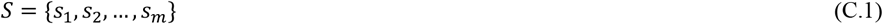

be a set of *m* distinct residues, and choose *s*_1_ ≡ *i* as the reference residue. Write

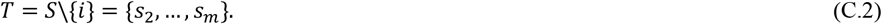

The *m*-th order effective distance is defined as the normalized Laplacian minor

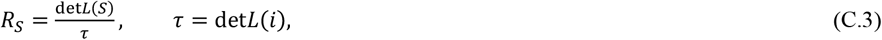

where *L*(*S*) denotes the matrix obtained from *L* by deleting the rows and columns indexed by the residues in *S*. For the one-residue reference set, the convention is *R*_{*i*}_ = 1.

We will show that

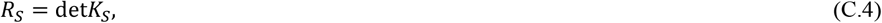

where *K*_*S*_ is the (*m* − 1) × (*m* − 1) symmetric matrix with entries

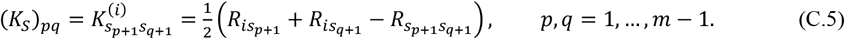

Thus the full hierarchy of higher-order effective distances can be evaluated using only the pairwise effective distances *R*_*ab*_.

#### C.2 Grounding at the reference residue

Deleting the row and column of the reference residue *i* gives the grounded Laplacian *L*(*i*). Since the graph is connected, *L*(*i*) is symmetric positive definite and invertible. Denote its inverse by

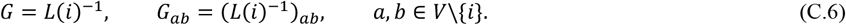

The pairwise effective distance has a direct expression in terms of *G*. Injecting unit current at residue *a* and extracting it at the grounded reference *i* gives

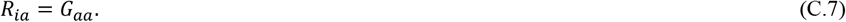

Injecting unit current at *a* and extracting it at *b* gives

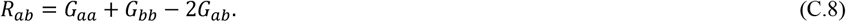

Substituting (C.7) and (C.8) into (C.5) yields

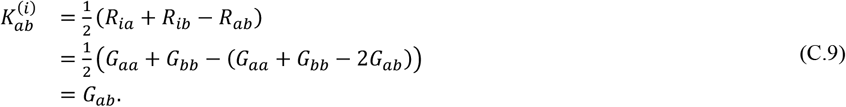

Therefore

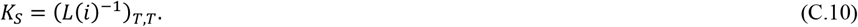

So *K*_*S*_ is not an arbitrary matrix of higher-order minors. It is the Gram matrix of the displacement vectors from the reference residue *i* to the non-reference residues in *S*.

#### C.3 The minor identity

We use Jacobi’s complementary-minor identity [20]. If *A* is invertible and *α* is an index set, then

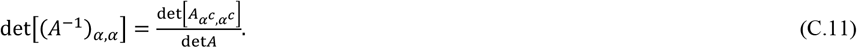

Apply (C.11) to *A* = *L*(*i*) and *α* = *T*. The index set of *L*(*i*) is *V*\{*i*}; deleting *T* from this grounded matrix removes the full set *S* = {*i*} ∪ *T* from the original Laplacian. Hence

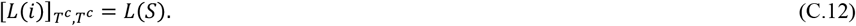

Using (C.10)-(C.12),

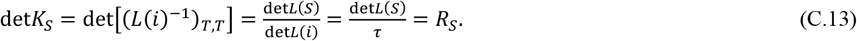

This proves the equivalence between the normalized Laplacian minor and the Gram determinant. The all-minors matrix-tree theorem gives the same result combinatorially, since det*L*(*S*) is a weighted sum over spanning forests rooted at the residues in *S* [14].

#### C.4 Geometric interpretation

By (C.10), *K*_*S*_ is a Gram matrix. The determinant of a Gram matrix equals the squared volume of the parallelotope spanned by the underlying vectors [20]. Therefore

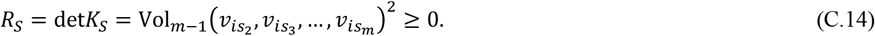

Thus the *m*-th order effective distance measures the squared collective volume generated by the communication directions from the reference residue *i* to the remaining residues in *S*. It vanishes precisely when these displacement vectors become linearly dependent, corresponding to collapse into a lower-dimensional communication subspace. The progression from second to fourth order has the geometric interpretation

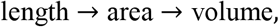

corresponding to *m* = 2,3,4. No simplex volume is used in the formulas above; the determinant directly represents the squared parallelotope volume.

#### C.5 Recovery of the lower orders Second order: *m* = 22

For *S* = {*i, j*}, *T* = {*j*} and

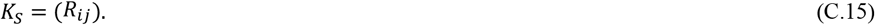

Therefore

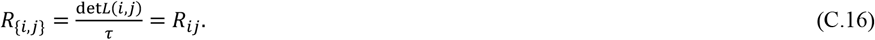

**Third order: *m*** = **33**

For *S* = {*i, j, k*}, *T* = {*j, k*} and

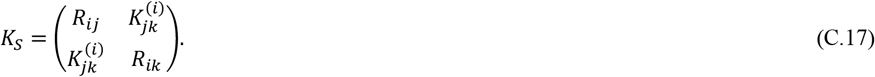

Thus

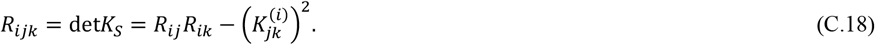

The third-order cooperation index is therefore

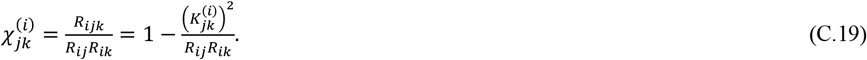

Equivalently, when the reference residue is clear, this may be written as *χ* _*ij K*_.

**Fourth order: *m*** = **44**

For *S* = {*i, j, k, l*}, *T* = {*j, k, l*} and the correct Gram matrix is

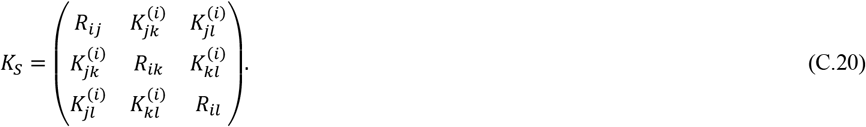

Therefore

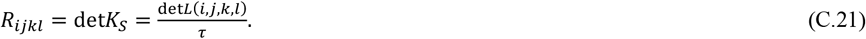

Expanding (C.20) gives

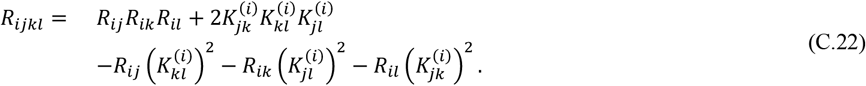

This fourth-order matrix is built from pairwise overlaps 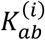, not from third-order effective distances. The quantities *R*_*ijk*_ and *R*_*ijkl*_ are determinants of Gram matrices; they are not themselves Gram-matrix entries.

#### C.6 Normalized all-order cooperation index and its range

We now define the normalized all-order cooperation index for the reference-labeled set *S* = {*i, s*_2_, …, *s*_*m*_}. Since the denominator depends on the chosen reference residue *i*, we write the index as 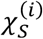:

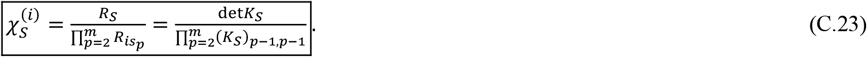

The product begins at *p* = 2 because *s*_1_ = *i* and *R*_*ii*_= 0. This notation avoids the ambiguity that would arise from writing a product over all residues in *S*.

The range of 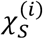 follows immediately from the Gram structure of *K*_*s*_. Since *K*_*s*_ is positive semidefinite,

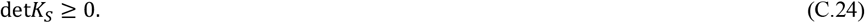

By Hadamard’s determinant inequality, the determinant of a positive semidefinite matrix is bounded above by the product of its diagonal entries [20]:

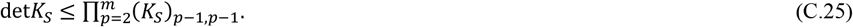

Using the diagonal identity

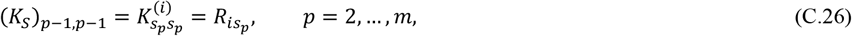

we obtain

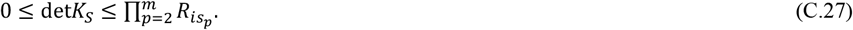

Dividing by the positive diagonal product gives

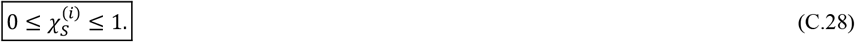

Thus every normalized all-order cooperation index lies in the closed interval [0,1]. The upper bound 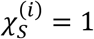 is reached when the displacement vectors from the reference residue to the other residues are mutually orthogonal. The lower bound 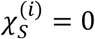 is reached when these displacement vectors are linearly dependent. In the language of the main text, this lower-bound limit corresponds to maximal cooperativity, whereas values near 1 correspond to independent communication channels.

The third-order index is the special case *m* = 3:

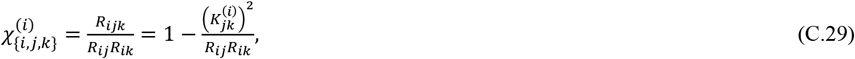

which recovers (C.19). The fourth-order index is

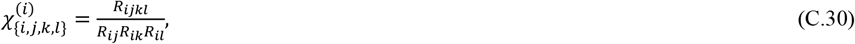

where *R*_*ijkl*_ is the determinant of the 3 × 3 Gram matrix in (C.20). Therefore the same Hadamard bound gives 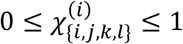.

#### C.7 Consequence: the Gram matrix is unique

Equations (C.3) and (C.8) determine the Gram matrix uniquely. Its off-diagonal entries are the pairwise overlaps 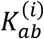, equivalently the entries *G*_*ab*_ of the grounded inverse *L*(*i*)^−1^. Higher-order quantities such as *R*_*ijkl*_ and *R*_*ijkl*_ are determinants of Gram matrices. They should not be inserted as off-diagonal entries of a new Gram matrix. Doing so defines a different object and breaks the identity *R*_*S*_ = det*K*_*S*_.

## Appendix D. Correspondence to Elastic Network Models

In the elastic network model, the thermal fluctuation inner product is:

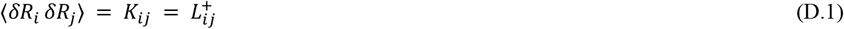

where *δR*_*i*_ is the displacement of residue i from its equilibrium position. This is the physical content of *K*: it is the correlation matrix of thermal fluctuations. Accordingly, the displacement vector *v*_*ij*_ given in the Methods section is the difference between the fluctuation vectors of residues j and i:

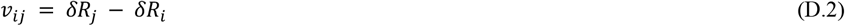

This is the relative displacement of residue j with respect to residue i. Its squared length in the fluctuation metric is:

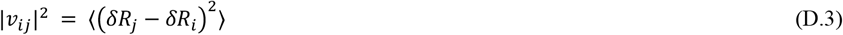

Expanding:

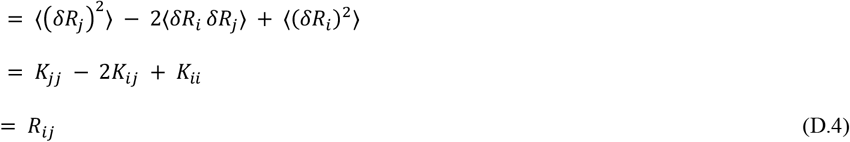

which is the effective distance. So the effective distance is simply the mean-squared relative displacement between residues i and j, a completely physical and intuitive quantity in the elastic network language.

### Now, the inner product of two displacement vectors

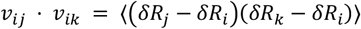

Expanding leads to Eq. B.7. Every term has a direct physical meaning in the elastic network language. 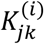 is the covariance of the relative displacements of j and k, both measured relative to the reference residue i, showing how correlated the fluctuations of residues j and k are if residue i is held fixed.

The angle:

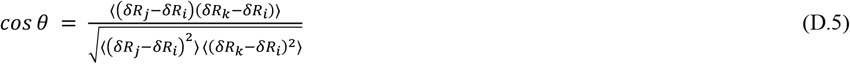

This is a normalized covariance, or Pearson-like normalized correlation, between the relative displacement of residue j from i and the relative displacement of residue k from i. By the Cauchy– Schwarz inequality,

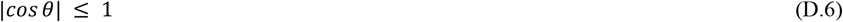

Therefore, the squared normalized overlap between the two anchored relative displacement directions is

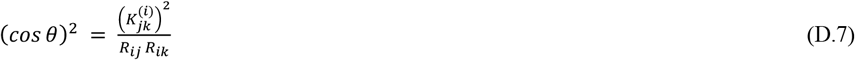

Using the cooperation-index definition from the Methods section, 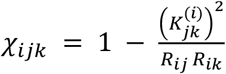, we obtain

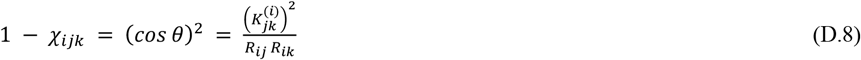

Thus, 1 − *χ*_*ijk*_ is the squared normalized overlap, or squared Pearson-like correlation, between the two relative fluctuation directions anchored at residue i.

The physical interpretation is therefore as follows. When *χ*_*ijk*_ = 0, the two anchored relative motions are perfectly correlated or anticorrelated, corresponding to maximal pathway overlap and strongest coupling. When *χ*_*ijk*_ = 1, the two anchored relative motions are orthogonal, corresponding to independent communication directions.

This interpretation preserves the sign convention used in the Results section: negative Δχ values indicate increased pathway overlap or increased coupling relative to the wild type, whereas values near zero indicate little change in the coupling state.

